# Xylazine exacerbates fentanyl-induced respiratory depression and prevents rescue by naloxone in mice

**DOI:** 10.1101/2025.06.18.660459

**Authors:** Joshua Watkins, Rachel Hahn, Michael Dempsey, Andrea G. Hohmann

## Abstract

Xylazine is a veterinary sedative and widespread adulterant of illicit opioids, where it is commonly combined with the highly potent synthetic µ opioid receptor (MOR) agonist fentanyl. Xylazine adulteration of fentanyl is associated with increased risk of lethal overdose and decreased efficacy of reversal by the MOR antagonist naloxone. Here we use whole body plethysmography in mice to show that xylazine produces profound respiratory depression at subanesthetic doses. Xylazine rapidly and dose-dependently suppressed minute ventilation, tidal volume, and respiratory frequency. These effects were dependent on α-2 adrenergic receptors and were fully blocked by coadministration of the α-2 adrenergic antagonist atipamezole. Atipamezole, administered alone, produced only modest reversal of fentanyl-induced respiratory depression. Xylazine, when combined with a dose of fentanyl with modest respiratory effects, suppressed breathing with greater efficacy than when administered alone. Strikingly, doses of naloxone sufficient to completely reverse fentanyl-induced respiratory depression were ineffective in reversing the respiratory suppression induced by xylazine-adulterated fentanyl. By contrast, combinations of naloxone with atipamezole rapidly and fully reversed the suppression of breathing induced by xylazine-adulterated fentanyl. Our results show that xylazine suppresses breathing via activation of α-2 receptors, an effect enhanced by coadministration with the MOR agonist fentanyl. Respiratory suppression inflicted by the mixture of xylazine and fentanyl resisted reversal by naloxone but was fully reversible by subsequent coadministration of both naloxone and atipamezole. These observations have profound implications for the current opioid epidemic.

## 1. Introduction

Xylazine, a relatively unregulated α-2 adrenergic receptor agonist and common veterinary sedative, is a widespread adulterant in the United States illicit opioid supply. Xylazine’s sedative effects are viewed as heightening and extending the rewarding effects of opioids [1–3]. Consequently, the mixture of xylazine with illicit opioid drugs – a substance commonly referred to as “tranq” – has become frequently and increasingly detected in lethal opioid overdose [4–11], as combinations of fentanyl and xylazine facilitate the use of lower doses of illicit opioids to produce subjectively similar effects at less risk and at lower cost [2]. Nearly all cases of overdose involving xylazine also involve the synthetic µ opioid receptor (MOR) agonist fentanyl [1, 11, 12] suggesting that xylazine-fentanyl coadministration may increase the risk of fatal overdose. These factors have driven the Office of National Drug Control Policy to declare xylazine-adulterated fentanyl an emerging threat to the United States, the first declaration of its kind in history [13].

While xylazine is a widely used in veterinary medicine – where it is commonly used as a sedative in combination with ketamine - it is not approved for human use due to risk of bradycardia and hypotension [14]. Consequently, the effects of xylazine alone or in concert with other drugs of abuse have rarely been investigated in a preclinical context [15]. Despite this, human use of xylazine is not rare. Illicit use of xylazine in North America was first detected in Puerto Rico in the year 2000 [16], and became a major adulterant of its heroin supply by 2006 [17]. By 2015 the combination of xylazine with illicit opioids was detected in the mainland United States [18], and xylazine use was detected in 48 states and over 10% of all overdoses involving fentanyl by 2022 [19, 20]. Use of xylazine-adulterated fentanyl has subsequently been detected across multiple Canadian provinces [21, 22] and across Europe, with the first case of xylazine-involved opioid overdose in the European Union identified in Italy in 2024 [23–25]. As suppression of respiratory drive is the primary cause of death in opioid overdose [26], we consequently hypothesized that xylazine may suppress breathing and exacerbate opioid-induced respiratory depression.

Experiments in rodent models support the idea that xylazine-fentanyl coadministration increases the lethality of opioid agonists [27, 28]. However, the mechanisms behind this increase in lethality are poorly understood. Xylazine induces severe bradycardia in humans [14, 17], which has largely precluded its study and use outside of veterinary settings. Conversely, the lethal effects of exogenous opioid drugs are well-studied, and largely attributable to the suppression of breathing [26]. Respiratory depression has been observed in human cases of xylazine poisoning [29–37], suggesting that xylazine-associated increases in opioid lethality may also be the result of suppressed breathing. This possibility is bolstered by observations that xylazine coadministration with fentanyl exacerbates opioid-induced hypoxia in the central nervous system [38].

Here we used whole body plethysmography in awake mice to demonstrate that xylazine induces severe respiratory depression at subanesthetic doses in both male and female mice. Furthermore, we show that the combination of xylazine and fentanyl produces respiratory depression more severe than that induced by fentanyl alone. We also demonstrate the pharmacological specificity of xylazine’s respiratory effects using the α-2 antagonist atipamezole. We found that the MOR antagonist naloxone was unable to reverse the respiratory effects of xylazine-adulterated fentanyl. Finally, we show that the combination of naloxone and atipamezole was able to reverse the respiratory suppressive effects of xylazine-adulterated fentanyl.

## 2. Materials and Methods

### 2.1 Animals

Subjects consisted of 164 male and 36 female C57BL/6J wild type mice (Jackson Laboratories; Bar Harbor, ME) aged 12 weeks at time of testing. Male mice weighed 30-40g, and female mice weighed 20-30g prior to testing. Mice were housed in a temperature- and humidity-controlled facility with ad libitum access to food and water on a 12:12h light/dark cycle. All procedures were approved by the Bloomington Institutional Animal Care and Use Committee of Indiana University.

### 2.2 Drugs

Fentanyl citrate and naloxone hydrochloride dihydrate were purchased from Sigma Aldrich (St. Louis, MO). Atipamezole hydrochloride was purchased from Tocris Bioscience (Bristol, UK). Xylazine injectable solution was purchased from Covetrus (Portland, ME). All drugs were administered intraperitoneally in a vehicle of 0.9% saline (Baxter Pharmaceuticals; Bloomington, IN) in a volume of 10 mL/kg.

### 2.3 Assessment of Respiratory Parameters

Whole body plethysmography (Data Sciences International; St. Paul, MN) was used to assess respiratory parameters of awake and unrestrained mice as described previously [39]. Respiratory metrics evaluated were minute ventilation (total volume of gas exchange, expressed as mL/minute), frequency (breaths per minute), and tidal volume (volume of individual breaths in mL).

Mice were habituated to recording chambers within 72 hours of the test day by placing them in recording chambers without experimental manipulation to minimize changes in breathing resulting from exploratory behavior or stress related to a novel environment. On the test day, mice received air mixed with 10% CO_2_ to standardize hypercapnic responses. Testing consisted of a 50-minute initial recording to establish baseline respiratory parameters for each subject, after which mice were briefly removed from test chambers and given an IP injection of drug or vehicle. Mice were then immediately returned to recording chambers, and recording was resumed. In experiments involving single injections, recordings continued for an additional 60 minutes without interruption. In experiments involving multiple injections, mice were briefly removed from the chambers after 15 minutes, given a second IP injection of either drug or vehicle, and then immediately returned to the chambers for an additional 45 minutes. Experiments were terminated immediately after the post-injection observation interval.

To establish the respiratory effects of xylazine, respiratory parameters were measured before and after a single i.p. injection of xylazine (0.1, 0.3, 1, 3, or 10 mg/kg) in male and female mice (n = 6-8 per group). Pharmacological specificity of xylazine’s respiratory effects and the role of α-2 adrenergic receptors was established by recording respiratory parameters before and after injection of xylazine (3 mg/kg) coadministered with the α-2 adrenergic receptor antagonist atipamezole (1 mg/kg) in male mice (n = 4-8 per group).

To assess the effects of xylazine-fentanyl coadministration, respiratory parameters were measured before and after a single i.p. injection of xylazine (0.1, 0.3, 1, 3, or 10 mg/kg) coadministered with a fixed dose of the synthetic MOR agonist fentanyl (0.2 mg/kg) in both male and female mice (n = 4-8 per group). The ability of naloxone (10 mg/kg) and atipamezole (3 mg/kg) to reverse the respiratory effects of fentanyl were evaluated by administering fentanyl (0.2 mg/kg) by i.p. injection to male mice (n = 4-6 per group), followed by a second i.p. injection of naloxone or atipamezole after fifteen minutes.

The efficacy of naloxone (10 mg/kg) in reversing the coadministration of fentanyl (0.2 mg/kg) and xylazine (3 mg/kg) was evaluated in male mice (n = 4-6 per group) by simultaneously administering xylazine and fentanyl followed by a second i.p. injection of naloxone after fifteen minutes. We similarly assessed the efficacy of combinations of naloxone and atipamezole in reversing xylazine fentanyl coadministration. We administered fentanyl (0.2 mg/kg) in combination with xylazine (3 mg/kg) by i.p. injection to male mice (n = 4-6 per group). After fifteen minutes, we administered a second i.p. injection containing naloxone (10 mg/kg) and atipamezole (1 mg/kg or 3 mg/kg).

### 2.4 Data Analysis

Statistical analyses were conducted using GraphPad Prism v.10.4 (Graphpad Software; Boston, MA). Plethysmography data were averaged into five-minute bins and represented as mean ± SEM. Two-way repeated measures ANOVA was used to assess the effects of pharmacological manipulations across time and to compare pre-injection baseline respiratory activity between experimental groups. Pre-injection and post-injection parameters were analyzed separately to ensure that differences in baseline respiratory activity did not impact interpretation of drug effects. Bonferroni’s *post hoc* test was used to assess whether pre-injection baseline parameters differed as a function of drug allocation, and whether antagonist conditions differed from active pharmacological manipulations or relevant controls. Dunnett’s *post hoc* test was used to compare the effects of active pharmacological manipulations to relevant controls. Dose response curves were generated by taking the difference between the final pre-injection bin and the bin twenty minutes after the first injection (a time point after maximal drug effect and before observed recovery for all drugs evaluated) and fitting a three-parameter logistic regression model using Graphpad Prism v.10.2.

Model-fitting was constrained to values below one hundred, as inhibition of respiratory parameters over one hundred percent would represent negative respiratory activity. Bottom values were constrained to zero in experiments in which the relevant comparator was a vehicle control, as the lowest tested doses in each experimental group did not significantly differ from vehicle controls for any tested metric. Bottom values were constrained to experimentally observed values in conditions where the relevant comparator was not a vehicle control (e.g., in evaluating the effects of xylazine-fentanyl coadministration, the bottom value of the model was set to the inhibition caused by the fixed fentanyl dose alone). Estimation of 95% confidence intervals for slope of respiratory suppression was calculated by simple linear regression in Graphpad Prism v.10.2.

## 3. Results

### 3.1 General results

Respiratory parameters generally showed a modest and expected decline across time, consistent with decreasing exploratory behavior and stress resulting from exposure to a novel environment as mice acclimated to recording chambers (p < 0.0234 for all conditions). No group differences in baseline minute ventilation were observed, and no group differences in other dependent measures assessing baseline respiratory activity for other metrics were observed except where specifically noted. No fatalities were observed following any experimental manipulations in any study.

### 3.2 Xylazine suppresses respiratory activity in both male and female mice

The α-2 adrenergic receptor agonist xylazine, administered systemically, markedly decreased minute ventilation in a dose- and time-dependent manner in both male (Drug: F_5,32_ = 137.7, p < 0.0001; Interaction: F_55,352_ = 9.169, p < 0.0001) and female (Drug: F_5,30_ = 109.4, p < 0.0001; Interaction: F_55,330_ = 8.210, p < 0.0001) mice (Fig. 1A-B). Xylazine produced similar decreases to minute ventilation in both sexes. *Post hoc* comparisons showed that all doses of xylazine 1 mg/kg and higher depressed minute ventilation in both male and female mice (1-10 mg/kg: p < 0.0001 for each sex). The EC_50_ (95% Confidence Interval; CI) for xylazine-induced suppression of minute ventilation was 1.133 (0.854 – 1.503) mg/kg in male mice and 1.005 (0.845 – 1.195) mg/kg in female mice (Fig. 1C).

**Figure 1.**
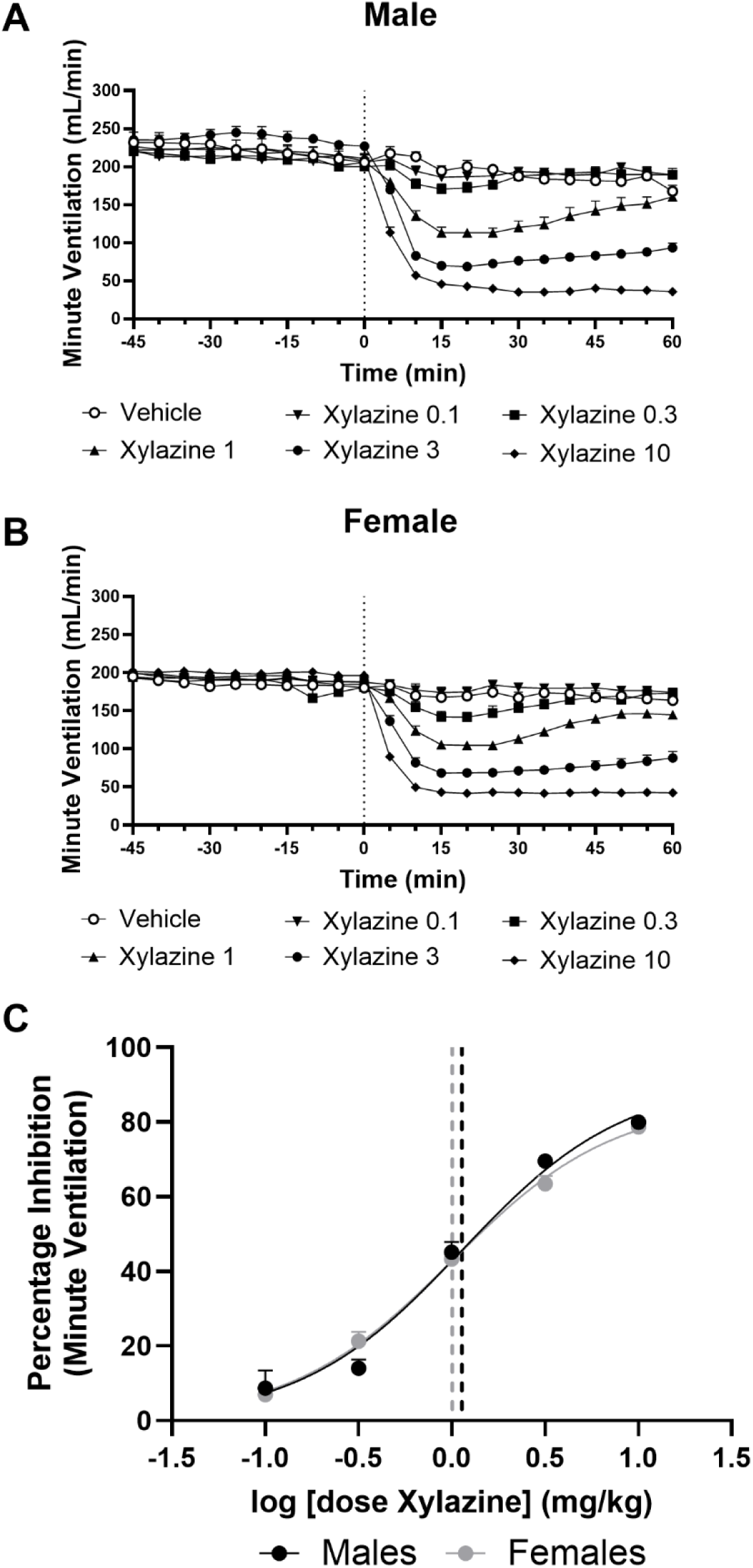
Xylazine reduces minute ventilation in mice of both sexes. Xylazine reduced minute ventilation in a dose- and time-dependent manner in both (A) male and (B) female mice. All doses 1 mg/kg and higher depressed minute ventilation in both male and female mice. (C) Dose response of xylazine’s suppression of minute ventilation in both male and female mice. EC_50_ (95% CI): Males: 1.133 (.0854 – 1.503) mg/kg i.p.; Females: 1.005 (0.8447 – 1.195) mg/kg i.p. Data were analyzed by two-way ANOVA followed by Dunnett’s *post hoc* test. Data are shown as Mean ± SEM where visible (n = 6-8 per group). Doses are shown as mg/kg, i.p.

Xylazine similarly suppressed respiratory frequency in both male (Drug: F_5, 32_ = 84.70, p < 0.0001; Interaction: F_55, 352_ = 6.596, p < 0.0001) and female (Drug: F_5, 30_ = 71.16, p < 0.0001; Interaction: F_55, 330_ = 2.826, p < 0.0001) mice (Fig. 2 A-B). Xylazine additionally suppressed tidal volume in both male (Drug: F_5.32_ = 71.79, p < 0.0001; Interaction: F_55,352_ = 15.01, p < 0.0001) and female (Drug: F _5,30_ = 63.65, p < 0.0001; Interaction: F _55,330_ = 10.21, p < 0.0001) mice (Fig. 2 C-D). All doses 1 mg/kg and higher suppressed both frequency (Males: 1-10 mg/kg, p < 0.0001; Females: 1 mg/kg, p = 0.0001; 3-10 mg/kg, p < 0.0001) and tidal volume (Males: 1 mg/kg, p = 0.0002; 3-10 mg/kg, p < 0.0001; Females: 1 mg/kg, p = 0.0002; 3-10 mg/kg, p < 0.0001) in both male and female mice.

**Figure 2.**
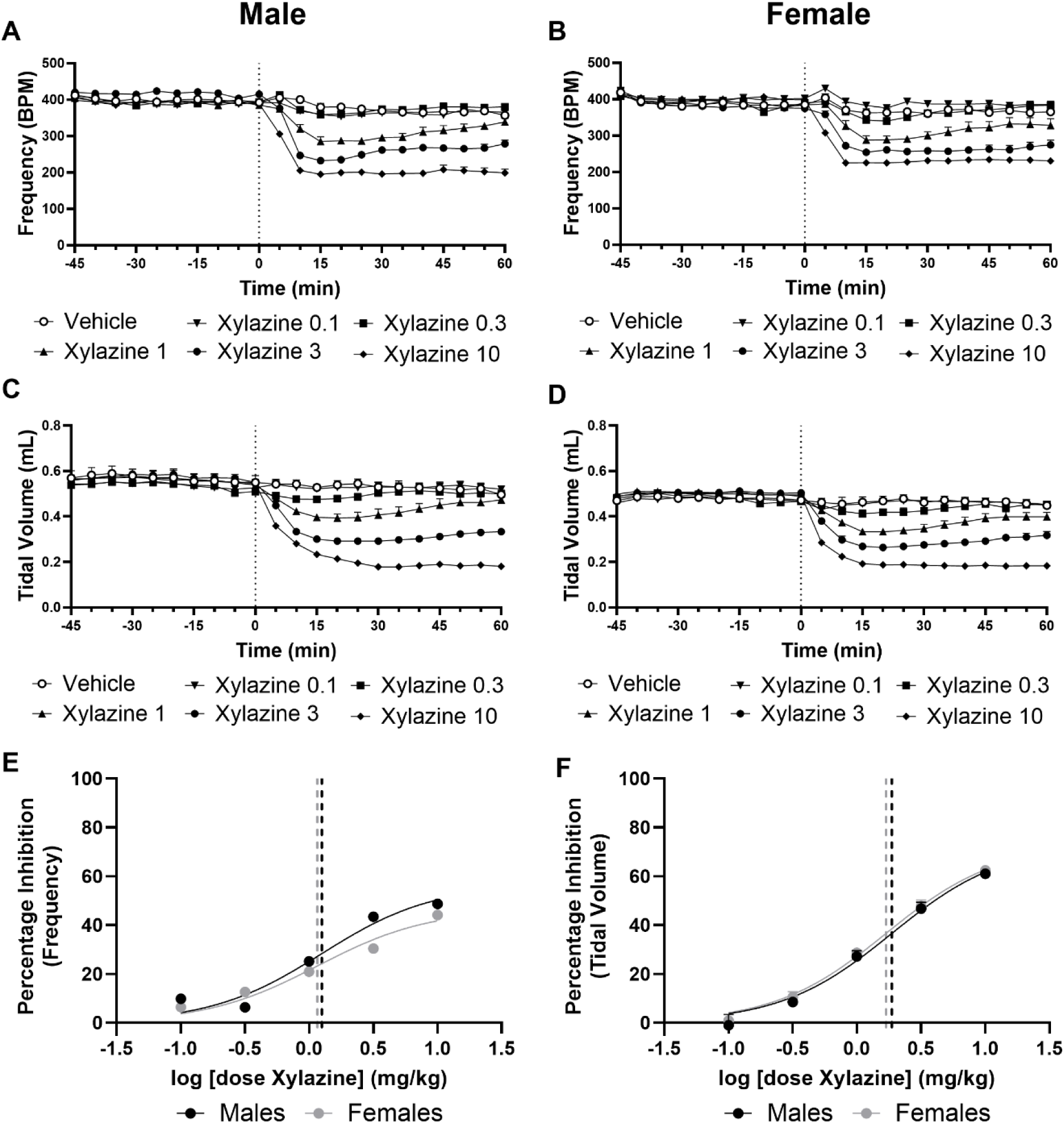
Xylazine reduced respiratory frequency in both (A) male and (B) female mice. Doses 1 mg/kg and higher suppressed frequency in both male and female mice. Xylazine similarly reduced tidal volume in a dose- and time-dependent manner in both (C) male and (D) female mice. All doses 1 mg/kg and higher depressed tidal volume in both male and female mice. (E-F) Dose response of xylazine’s suppression of (E) frequency and (F) tidal volume in mice of both sexes. EC_50_ (95% CI) tidal volume: Males: 1.876 (1.284 – 2.764) mg/kg i.p.; Females: 1.686 (1.333 – 2.139) mg/kg i.p. EC_50_ (95% CI) frequency: Males: 1.255 (0.090 – 1.747) mg/kg i.p.; Females 1.159 (0.766 – 1.752) mg/kg i.p. Data were analyzed by two-way ANOVA followed by Dunnett’s *post hoc* test. Data are shown as Mean ± SEM where visible (n = 6-8 per group). Doses are shown as mg/kg, i.p.

The EC_50_ for xylazine-induced suppression of respiratory frequency was 1.255 (0.900 – 1.747) mg/kg in male mice and 1.159 (0.766 – 1.752) mg/kg in female mice (Fig. 2E). The EC_50_ for xylazine-induced suppression of tidal volume was 1.874 (1.284 – 2.764) mg/kg in male mice and 1.686 (1.333 – 2.139) mg/kg in female mice (Fig. 2F). In all analyses where dose response curves were generated, the 95% confidence intervals for metrics impacted by xylazine in male and female mice overlapped.

### 3.3 Xylazine-induced suppression of respiratory activity is completely prevented by coadministration of the α-2 adrenergic receptor antagonist atipamezole

The α-2 adrenergic receptor antagonist atipamezole (1 mg/kg, i.p.) completely blocked the respiratory suppressive effects of a coadministered dose of xylazine (3 mg/kg, i.p.) on minute ventilation (Drug: F_3, 20_ = 52.58, p < 0.0001; Interaction: F_33, 220_ = 7.187, p < 0.0001; Fig. 3A), respiratory frequency (Drug F_3, 20_ = 29.89, p < 0.0001; Interaction F_33, 220_ = 8.480, p < 0.0001; Fig. 3B), and tidal volume (Drug F_3, 20_ = 36.63, p < 0.0001; Interaction F_33, 220_ = 8.450, p < 0.0001; Fig 3C). Respiratory parameters were lower in groups receiving xylazine (3 mg/kg i.p.) alone compared to all other groups (p < 0.0001 for all metrics). Respiratory parameters of mice injected with only vehicle did not differ from those injected with both xylazine and atipamezole (p > 0.9999 for all metrics; Fig. 3A-C) or atipamezole alone (p > 0.9999 for all metrics; Fig. 3A-C).

**Figure 3.**
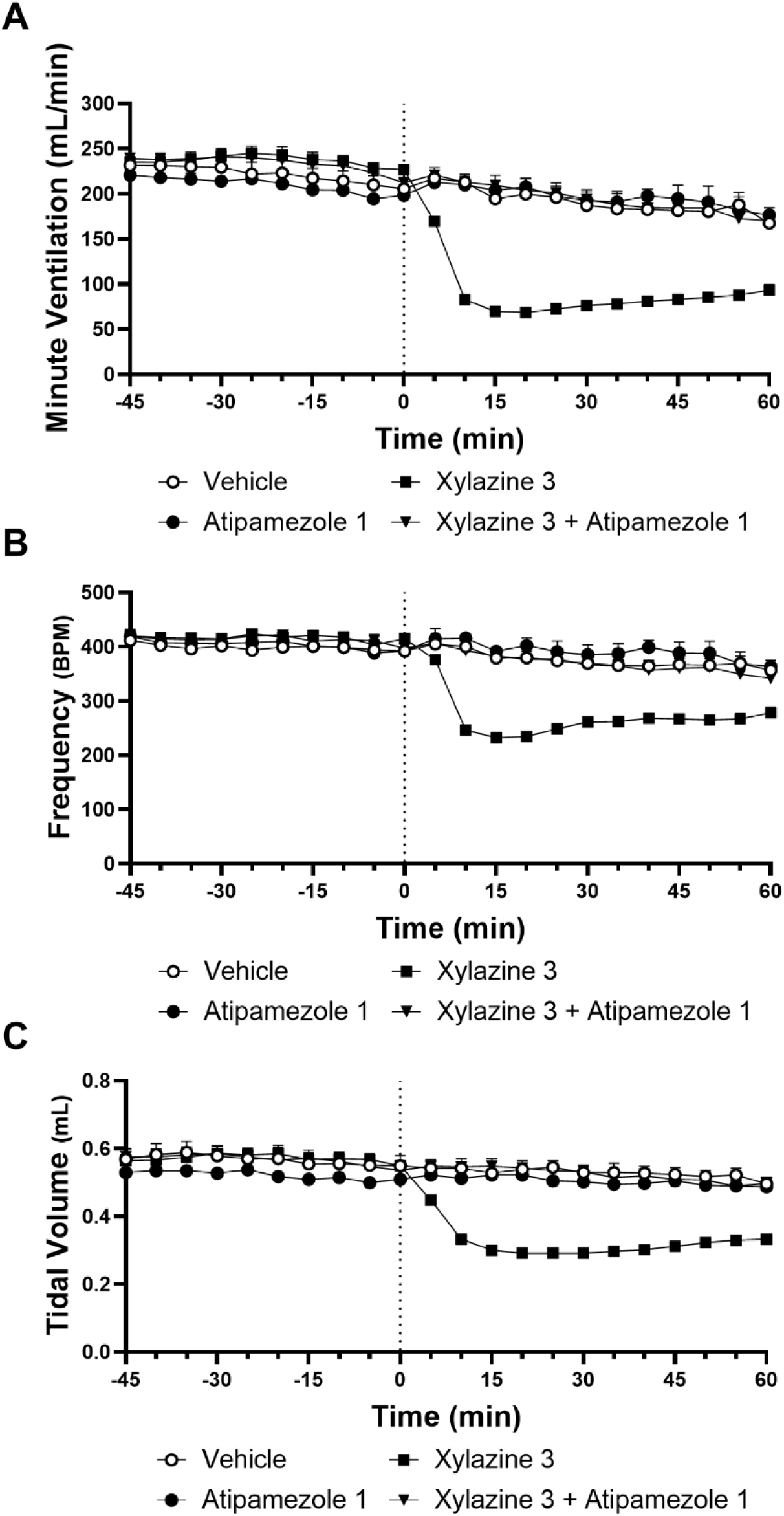
The α-2 antagonist atipamezole prevents the respiratory effects of xylazine. Coadministration of xylazine and atipamezole completely prevents reduction of (A) minute ventilation, (B) frequency, and (C) tidal volume in male mice. Data were analyzed by two-way ANOVA followed by Bonferroni’s *post hoc* test. Data are shown as Mean ± SEM where visible (n = 6-8 per group). Doses are shown as mg/kg, i.p.

### 3.4 Fentanyl-xylazine coadministration produces stronger suppression of respiratory activity than fentanyl alone

Fentanyl (0.2 mg/kg, i.p.) suppressed minute ventilation in the presence and absence of xylazine (0.1 – 10 mg/kg i.p.) and these effects were both dose- and time-dependent (Drug: F_6, 25_ = 40.80, p < 0.0001; Interaction: F_66, 275_ = 3.603, p < 0.0001; p = 0.0092 vs. vehicle) (Fig. 4A). Fentanyl (0.2 mg/kg, i.p.) suppressed minute ventilation relative to vehicle (p = 0.0092) whereas the same fentanyl dose coadministered with xylazine (1-10 mg/kg, i.p.) produced greater suppression of minute ventilation than fentanyl alone (1 mg/kg: p = 0.0440; 3-10 mg/kg: p < 0.0001; Fig. 4A). Coadministration of lower xylazine doses (0.1 -0.3 mg/kg) did not enhance fentanyl-induced suppression of minute ventilation relative to fentanyl alone (p > 0.999; Fig. 4A).

**Figure 4.**
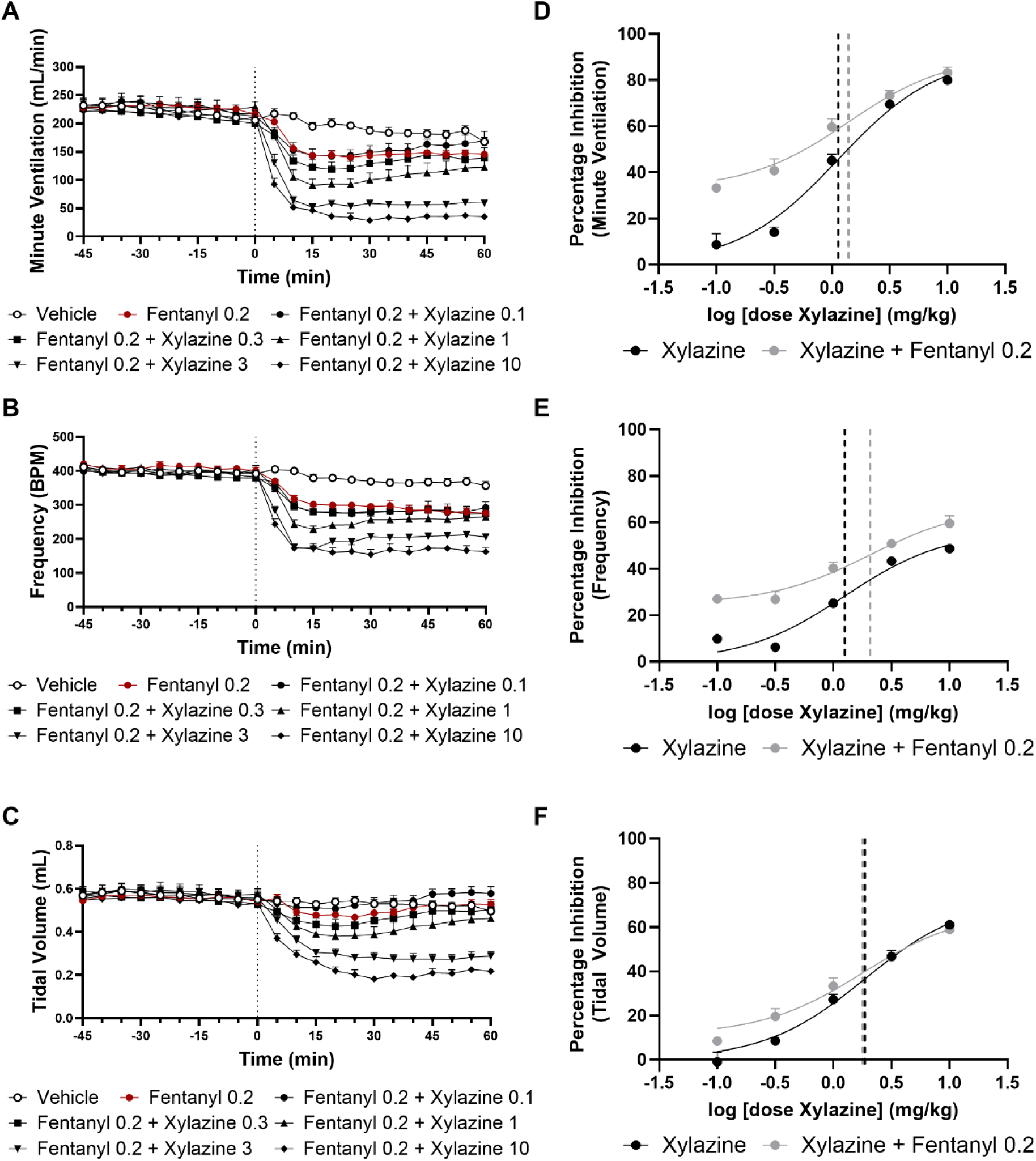
The combination of xylazine with a dose of the synthetic MOR agonist fentanyl sufficient to produce modest respiratory depression produced greater suppression of (A) minute ventilation, (B) frequency, and (C) tidal volume than fentanyl alone. Minute ventilation was suppressed to a greater degree by xylazine doses 1 mg/kg and above, frequency by doses 3 mg/kg and above, and tidal volume by doses 3 mg/kg and above. (D-F) Dose response of xylazine and xylazine-fentanyl combination in male mice. EC_50_ (95% CI) xylazine: minute ventilation: 1.133 (0.8542 – 1.503) mg/kg i.p.; frequency: 1.255 (0.8999 – 1.747) mg/kg i.p.; tidal volume: 1.874 (1.284 – 2.764) mg/kg, i.p. EC_50_ (95% CI) xylazine-fentanyl coadministration minute ventilation: 1.393 (0.8552 – 2.292) mg/kg i.p.; frequency: 2.080 (1.120 to 4.010) mg/kg, i.p.; tidal volume: 1.787 (1.032 to 3.170) mg/kg, i.p. Data were analyzed by two-way ANOVA followed by Bonferroni’s *post hoc* test. Data are shown as Mean ± SEM where visible (n = 6-8 per group). Doses are shown as mg/kg, i.p.

Fentanyl (0.2 mg/kg, i.p.) also suppressed respiratory frequency (Drug: F_6, 25_ = 48.07, p < 0.0001; Interaction: F_66, 275_ = 3.123, p < 0.0001; p < 0.0001 vs. vehicle; Fig. 4B) in male mice. Doses of xylazine 3 mg/kg and above produced greater suppression of frequency when coadministered with fentanyl than fentanyl produced alone (3-10 mg/kg, p < 0.0001; Fig. 4B). Fentanyl (0.2 mg/kg, i.p.) did not reliably suppress tidal volume (Drug: F_6, 25_ = 23.99, p < 0.0001; Interaction: F_66, 275_ = 8.281, p < 0.0001; p > 0.9999 vs. vehicle; Fig. 4C) in male mice. Doses of xylazine 3 mg/kg and above produced greater suppression of respiratory frequency when coadministered with fentanyl than fentanyl produced alone (3-10 mg/kg, p < 0.0001; Fig. 4C).

The EC_50_ dose for suppression of minute ventilation by xylazine in combination with fentanyl (0.2 mg/kg, i.p.) was 1.393 (0.855 – 2.292) mg/kg with a slope of 26.48 (22.38 – 30.57). The EC_50_ dose for xylazine alone was 1.133 (0.854 – 1.503) mg/kg with a slope of 39.59 (34.90 – 44.29) (Fig. 4D). The EC_50_ dose for suppression of frequency by xylazine in combination with fentanyl (0.2 mg/kg, i.p.) was 2.080 (1.120 to 4.010) mg/kg with a slope of 17.83 (14.09 – 21.58). The EC_50_ dose for xylazine alone was 1.255 (0.8999 – 1.747) mg/kg with a slope of 22.95 (19.45 – 26.44) (Fig. 4E). The EC_50_ dose for suppression of tidal volume by xylazine in combination with fentanyl (0.2 mg/kg, i.p.) was 1.787 (1.032 – 3.170) mg/kg with a slope of 25.26 (21.98 – 29.14). The EC_50_ dose for xylazine alone was 1.874 (1.284 – 2.764) mg/kg with a slope of 32.45 (28.84 –36.06) (Fig. 4F).

### 3.5 Both naloxone and atipamezole reverse fentanyl-induced respiratory depression

The MOR antagonist naloxone (10 mg/kg, i.p.), administered 15 minutes after fentanyl (0.2 mg/kg, i.p.), completely reversed the opioid-induced suppression of minute ventilation overall and in a time-dependent manner (Drug: F_2, 14_ = 10.11, p = 0.0019; Interaction: F_16, 112_ = 6.627, p < 0.0001; Fig. 5A). Minute volume was lower overall in groups that received fentanyl followed by vehicle treatment, compared to groups receiving either fentanyl followed by naloxone rescue (p = 0.0217) or vehicle followed by vehicle (p = 0.0021). Naloxone rescue completely reversed fentanyl-induced suppression of minute ventilation (p = 0.612 vs. vehicle followed by vehicle).

**Figure 5.**
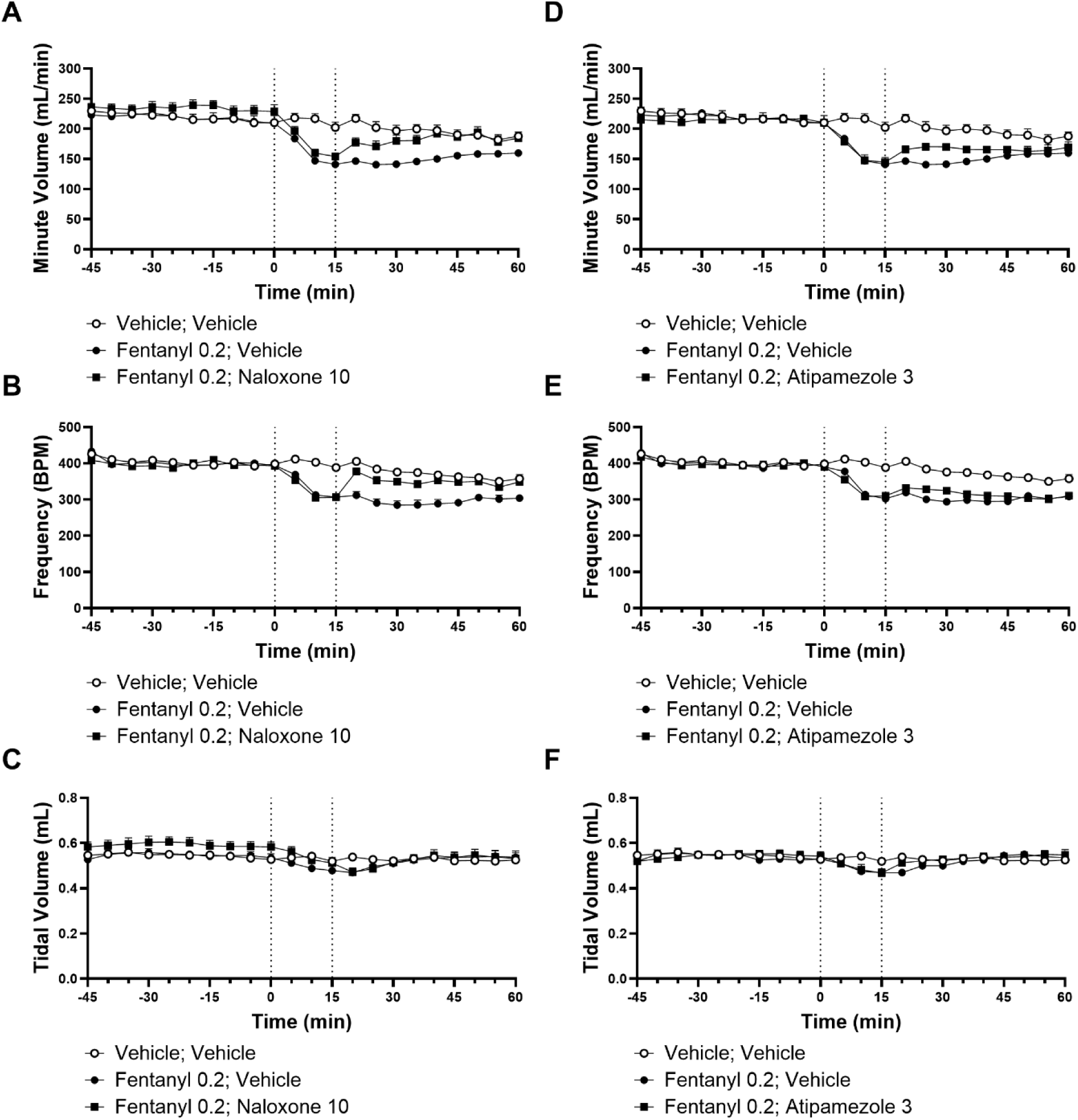
The MOR antagonist naloxone reverses the respiratory effects of fentanyl. Administration of naloxone fifteen minutes after administration of fentanyl completely reversed fentanyl-induced reduction of (A) minute ventilation and (B) frequency. Neither metric differed from mice exposed to vehicle. (C) Tidal volume was not altered by this dose of fentanyl, and no group differed from vehicle controls. Atipamezole (3 mg/kg, i.p.), administered fifteen minutes after fentanyl (0.2 mg/kg, i.p.) produced a modest reversal of fentanyl-induced suppression of (D) minute ventilation and (E) frequency. This reversal was limited to timepoints 10 and 15 minutes after the injection of atipamezole. This dose of fentanyl did not depress tidal volume (F). Data were analyzed by two-way ANOVA followed by Bonferroni’s *post hoc* test. Data are shown as Mean ± SEM where visible (n = 4-6 per group). Doses are shown as mg/kg, i.p.

Similarly, fentanyl (0.2 mg/kg i.p.) followed by vehicle challenge reduced respiratory frequency in a naloxone-reversible manner (Drug: F_2, 14_ = 24.43, p < 0.0001; Interaction: F_16,112_ = 2.951, p = 0.0004; Fig. 5B). Respiratory frequency was lower in groups that received fentanyl followed by vehicle treatment compared to groups receiving fentanyl followed by naloxone rescue (p = 0.0005) or vehicle followed by vehicle (p < 0.0001).

Naloxone rescue completely reversed fentanyl-induced suppression of respiratory frequency (p = 0.274 vs. vehicle followed by vehicle). This dose of fentanyl did not suppress tidal volume overall although the interaction was significant (Drug: F_2,14_ = 0.0281, p = 0.9724; Interaction: F_16,112_ = 5.221, p < 0.0001; Fig. 5C). *Post hoc* comparisons failed to reveal any differences between groups at any post-injection timepoint (p ≥ 0.1043).

The α-2 adrenergic receptor antagonist atipamezole (3 mg/kg, i.p.) partially reversed the fentanyl-induced suppression of minute ventilation and these effects were time dependent (Drug: F_2,12_ = 13.63, p = 0.0008; Interaction: F_16,96_ = 7.019, p < 0.0001; Fig. 5D). Minute ventilation was lower overall in groups receiving either fentanyl followed by vehicle (p = 0.0007) or fentanyl followed by atipamezole (p = 0.0234) compared to the vehicle-vehicle group. Atipamezole partially and transiently reversed fentanyl-induced suppression of minute ventilation at a subset of post-injection timepoints following injection of the α-2 adrenergic receptor antagonist (5 min: p = 0.0230; 10 min: p = 0.0440 vs. fentanyl followed by vehicle). Fentanyl-vehicle reduced minute ventilation compared to vehicle-vehicle treatment over fifty minutes after initial injection of fentanyl (p < 0.0209) and again at sixty minutes (p = 0.0178). Fentanyl-atipamezole treatment showed lower minute ventilation compared to vehicle-vehicle treatment for 45 minutes following initial injection of fentanyl (p < 0.0353) and again at 50 minutes (p = 0.0360) but was no longer present at 55 or 60 minutes (p ≥ 0.2307) post-injection.

Atipamezole (3 mg/kg, i.p.) challenge, administered fifteen minutes after fentanyl (0.2 mg/kg, i.p.), also produced a modest but reliable blunting of fentanyl-induced suppression of respiratory frequency and these effects were time-dependent (Drug: F_2,12_ = 27.97, p < 0.0001; Interaction: F_16,96_ = 3.649, p < 0.0001; Fig. 5E). Respiratory frequency was lower across the observation interval in both the fentanyl-vehicle group (p < 0.0001) and the fentanyl-atipamezole group (p = 0.0010) compared to the vehicle-vehicle group and the effects of fentanyl-vehicle did not differ from fentanyl-atipamezole treatment overall (p = 0.3206). Respiratory frequency was transiently higher in the fentanyl-atipamezole group compared to the fentanyl-vehicle group at a subset of post-injection timepoints (5 min: p = 0.0163; 10 min: p = 0.0117). By contrast, respiratory frequency was reduced in both the fentanyl-vehicle group (p < 0.0001) and the fentanyl-atipamezole group (p ≤ 0.0002) compared to the vehicle-vehicle group at all post-injection timepoints.

In fentanyl-treated groups that were subsequently challenged with either vehicle or atipamezole, tidal volume was not altered overall but the interaction across time was significant (Drug: F_2,12_ = 0.0976, p = 0.9077; Interaction F_16,96_ = 5.006, p < 0.0001; Fig. 5F). However, *post hoc* comparisons failed to reveal any differences between any groups at any post-injection time point (p ≥ 0.0712 fentanyl-vehicle vs. vehicle-vehicle; p > 0.9999 fentanyl-atipamezole vs. vehicle-vehicle; p ≥ 0.5446 fentanyl-vehicle vs. fentanyl-atipamezole).

### 3.6 Combinations of naloxone and atipamezole – but not naloxone alone – reverse respiratory suppression by xylazine-fentanyl coadministration

Fentanyl (0.2 mg/kg, i.p.) coadministered with xylazine (3 mg/kg, i.p.) suppressed minute ventilation in a manner that was preferentially reversed by the combination of naloxone (10 mg/kg., i.p.) and atipamezole (1-3 mg/kg, i.p.) but not by naloxone (10 mg/kg, i.p.) alone (Drug: F_4,20_ = 39.96, p < 0.0001; Interaction: F_32,160_ = 8.675, p < 0.0001). *Post hoc* comparisons revealed that coadministered fentanyl and xylazine reduced minute ventilation overall compared to either vehicle-vehicle treatment (p < 0.0001), or the same concentrations of xylazine-adulterated fentanyl that were subsequently challenged with a combination of naloxone (10 mg/kg, i.p.) and atipamezole (1 or 3 mg/kg, i.p.; p < 0.0001 for each comparison).

The combination of naloxone with the high (p = 0.5081) but not the low (p = 0.0044) dose of atipamezole reversed reductions in minute ventilation induced by xylazine-adulterated fentanyl to levels exhibited by mice receiving only vehicle (Fig. 6A). Notably, xylazine-adulterated fentanyl reduced minute ventilation similarly to that observed in groups challenged only with naloxone or with vehicle (p = 0.2755), showing the failure of naloxone rescue (Fig. 6A).

**Figure 6.**
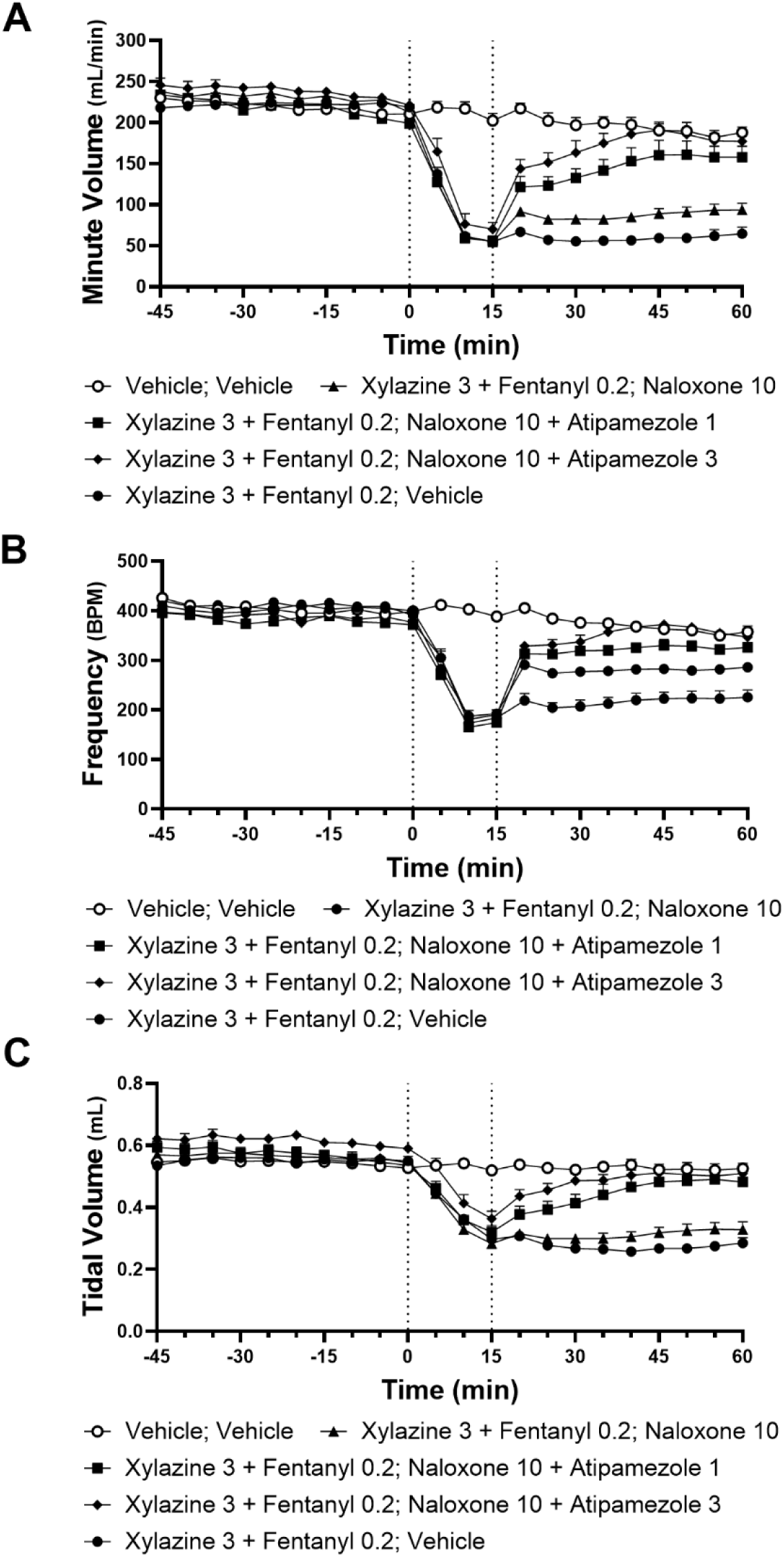
Naloxone failed to reverse the respiratory effects of xylazine-fentanyl coadministration. Naloxone (10 mg/kg, i.p.), administered fifteen minutes after the injection of a combination of xylazine (3 mg/kg, i.p.) and fentanyl (0.2 mg/kg, i.p.), did not reverse suppression of minute ventilation (A). Naloxone produced a modest but consistent increase in frequency (B) but failed to return to levels observed in vehicle controls. Naloxone did not reverse suppression of tidal volume (C). The combination of atipamezole and naloxone successfully reversed the respiratory effects of xylazine-fentanyl coadministration. Naloxone (10 mg/kg) and atipamezole (1 mg/kg) reversed the suppression of xylazine-fentanyl coadministration on (A) minute ventilation, (B) frequency and (C) tidal volume. Minute ventilation and frequency remained suppressed in comparison to vehicle controls, while tidal volume did not differ in comparison to mice receiving only vehicle. The combination of naloxone (10 mg/kg) and atipamezole (3 mg/kg) completely reversed the effects of xylazine-fentanyl coadministration at the same doses, reversing the suppression of (A) minute ventilation, (B) frequency and (C) tidal volume. All metrics did not differ from vehicle controls. Data were analyzed by two-way ANOVA followed by Bonferroni’s *post hoc* test. Data are shown as Mean ± SEM where visible (n = 4-6 per group). Doses are shown as mg/kg, i.p. Modest but consistent differences in baseline values for tidal volume (Drug: F_4,20_ = 5.500, p = 0.0037; Interaction: F_36,180_ = 1.212, p = 0.2069) and respiratory frequency (Drug: F_4,20_ = 5.401, p = 0.0041; Interaction: F_36,180_ = 1.720, p = 0.0082) were detected in experimental animals. *Post hoc* tests revealed differences in three conditions with respect to baseline measures of tidal volume. Differences were observed in baseline responsiveness between mice receiving vehicle followed by vehicle and mice receiving xylazine (3 mg/kg) and fentanyl (0.2 mg/kg) followed by naloxone (10 mg/kg) and atipamezole (3 mg/kg) (p = 0.0039), mice receiving xylazine (3 mg/kg) and fentanyl (0.2 mg/kg) followed by naloxone (10 mg/kg) and mice receiving xylazine (3 mg/kg) and fentanyl (0.2 mg/kg) followed by naloxone (10 mg/kg) and atipamezole (3 mg/kg) (p = 0.0374), and animals receiving animals receiving xylazine (3 mg/kg) and fentanyl (0.2 mg/kg) followed by naloxone (10 mg/kg) and atipamezole and animals receiving animals receiving xylazine (3 mg/kg) and fentanyl (0.2 mg/kg) followed by vehicle (p = 0.0165). *Post hoc* tests revealed differences in one condition with respect to baseline measures of respiratory frequency. Differences in baseline responsiveness in respiratory frequency were observed between mice receiving xylazine (3 mg/kg) and fentanyl (0.2 mg/kg) followed by naloxone (10 mg/kg) and mice receiving xylazine (3 mg/kg) and fentanyl (0.2 mg/kg) followed by naloxone (10 mg/kg) and atipamezole (1 mg/kg) (p = 0.0518). No differences were observed in baseline measures of overall respiratory activity as measured by minute ventilation (p = 0.1449).

Fentanyl (0.2 mg/kg, i.p.) coadministered with xylazine (3 mg/kg, i.p.) suppressed respiratory frequency in a manner that was preferentially reversed by the combination of naloxone (10 mg/kg., i.p.) and atipamezole (1-3 mg/kg, i.p.) but only partially reversed by naloxone (10 mg/kg, i.p.) rescue alone (Drug: F_4, 20_ = 55.65, p < 0.0001; Interaction: F_32, 160_ = 4.459, p < 0.0001: Fig. 6B). Xylazine-fentanyl coadministration reduced respiratory frequency relative to vehicle-vehicle treatment (p < 0.0001) or groups receiving the same concentration of xylazine-adulterated fentanyl that were subsequently challenged either with naloxone only, or naloxone in combination with either low (1 mg/kg, i.p.) or high (3 mg/kg, i.p.) dose atipamezole (p < 0.0001 for each comparison).

Naloxone (10 mg/kg i.p.) challenge increased respiratory frequency in groups receiving xylazine-adulterated fentanyl (p < 0.0001) but failed to restore levels of respiratory frequency to that observed in vehicle controls (p < 0.0001). The combination of naloxone with the high (p = 0.7037) but not the low (p = 0.0018) dose of atipamezole completely reversed the reductions in respiratory frequency induced by xylazine-adulterated fentanyl to levels observed in mice receiving only vehicle (Fig. 6B).

Fentanyl (0.2 mg/kg, i.p.) coadministered with xylazine (3 mg/kg, i.p.) suppressed tidal volume in a manner that was preferentially reversed by the combination of naloxone (10 mg/kg., i.p.) and atipamezole (1-3 mg/kg, i.p.) but not by naloxone (10 mg/kg, i.p.) rescue alone (Drug: F_4,20_ = 28.99, p < 0.0001; Interaction: F_32,160_ = 7.203, p < 0.0001; Fig. 6C). *Post hoc* comparisons revealed that xylazine-fentanyl coadministration reduced tidal volume overall relative to either vehicle-vehicle treatment (p < 0.0001), or groups receiving the same concentrations of xylazine-adulterated fentanyl that were subsequently challenged with a combination of atipamezole (1 or 3 mg/kg, i.p.) and naloxone (p < 0.0001 for each comparison).

The combination of naloxone with either the high (p > 0.9999) or the low (p = 0.0791) dose of atipamezole reversed reductions in minute ventilation induced by xylazine-adulterated fentanyl to levels observed in groups receiving only vehicle (Fig. 6C). Xylazine-adulterated fentanyl reduced tidal volume similarly in groups were challenged with vehicle or naloxone only (p > 0.9999), showing failure of naloxone rescue (Fig. 6C).

Administration of vehicle followed by naloxone (10 mg/kg, i.p.) or naloxone combined with atipamezole (3 mg/kg, i.p.) did not impact minute ventilation in comparison to mice receiving only vehicle (Drug: F_2,10_ = 0.0676, p = 0.9351; Interaction: F_16,80_ = 0.4175, p = 0.9740; Fig. 7A). Similarly, neither administration of naloxone alone or combined with atipamezole affected respiratory frequency (Drug: F_2,10_ = 0.0590, p = 0.5727; Interaction: F_16,80_ = 1.835, p = 0.0728; Fig. 7B). Measures of tidal volume did not differ between groups with respect to drug (Drug: F_2,10_ = 0.0117, p = 0.9884; Interaction: F_16,80_ = 1.835, p = 0.0404; Fig. 7C) although the interaction was significant. *Post hoc* comparisons revealed no reliable differences between groups at any timepoint (p > 0.9999 all timepoints).

**Figure 7.**
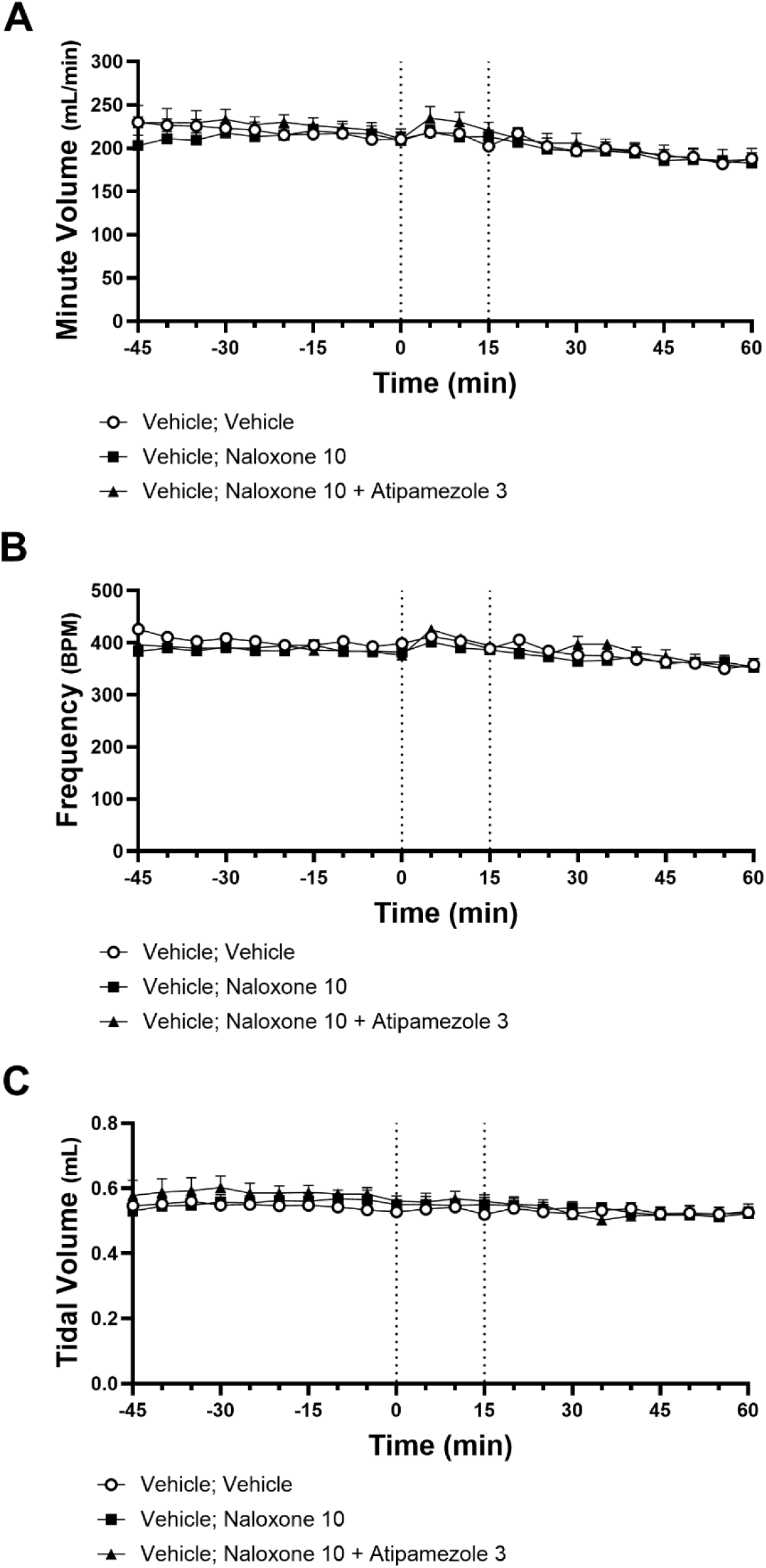
The MOR antagonist naloxone (10 mg/kg), alone or in combination with the α2 antagonist atipamezole (3 mg/kg), did not alter minute ventilation (A), respiratory frequency (B), or tidal volume (C) when administered fifteen minutes after vehicle challenge. Data were analyzed by two-way ANOVA followed by Bonferroni’s *post hoc* test. Data are shown as Mean ± SEM where visible (n = 4-6 per group). Doses are shown as mg/kg. i.p.

## 4. Discussion

Forensic drug reports increasingly link xylazine, a common veterinary anesthetic, to lethal fentanyl overdose in humans [4–11, 40]. Here we examined the effects of xylazine on breathing in the presence and absence of fentanyl and document failure of naloxone to reverse the respiratory suppressive effects of xylazine-adulterated fentanyl. We employed whole body plethysmography in adult C57Bl/6 mice of both sexes to examine how subanesthetic doses of xylazine impact respiration. Xylazine, administered alone at subanesthetic doses, induced respiratory depression by suppressing both respiratory frequency and tidal volume. Furthermore, xylazine in combination with the highly potent MOR agonist fentanyl enhanced the efficacy of low doses of fentanyl to suppress minute ventilation, tidal volume, and respiratory frequency. Xylazine’s respiratory effects were mediated by α-2 adrenergic receptors and were preventable by coadministration of the α-2 adrenergic receptor antagonist atipamezole. Moreover, atipamezole partially reversed fentanyl-induced respiratory depression with limited efficacy. Most relevant to clinical cases of human overdose, we show that the MOR antagonist naloxone does not effectively reverse respiratory depression induced by xylazine-fentanyl coadministration, while mixtures of naloxone and atipamezole, administered concurrently, completely reverse the respiratory suppressive effects of xylazine-fentanyl coadministration when administered fifteen minutes after xylazine-fentanyl coadministration.

Xylazine, administered alone at subanesthetic doses [41], suppressed respiratory activity in both male and female mice in a dose-dependent manner. Systemic doses of xylazine depressed both respiratory frequency and tidal volume, resulting in rapid, persistent, and profound depression of breathing as measured by minute ventilation. These effects were observed at low doses with human relevance [42]. Furthermore, the combination of xylazine with fentanyl produced more extreme suppression of breathing than fentanyl alone when combined with doses of xylazine that intrinsically suppressed breathing, mirroring the use of xylazine as an adulterant to illicit synthetic opioids.

The respiratory effects of xylazine were not sex-specific. In all instances where dose response curves were generated, the confidence intervals for metrics impacted by xylazine and xylazine-fentanyl coadministration overlapped, suggesting that drug effect did not differ between sexes. This observation aligns with previous reports that xylazine-induced lethality is generally similar between sexes [43].

Xylazine’s depression of breathing was dependent on α-2 adrenergic receptors. The sedative effects of xylazine are primarily attributed to its activity at the α-2 adrenergic receptor, at which xylazine functions as a nonselective agonist of all receptor subtypes [44]. α-2 receptors are inhibitory (Gi/Go) G protein-coupled receptors whose endogenous ligands are epinephrine and norepinephrine [45–47]. These receptors are expressed in both the central nervous system and the periphery, including in the brainstem, where breathing is primarily controlled [47–50]. The α-2 adrenergic receptor antagonist atipamezole completely prevented the effects of xylazine on breathing, and combinations of naloxone and atipamezole reversed the respiratory effects of xylazine-fentanyl coadministration in a subset of tested doses.

Suppression of breathing is observed in cases of human xylazine overdose, cases in which mechanical ventilation was required to counter confirmed xylazine overdoses, or in veterinary use of xylazine-ketamine anesthesia [29–37, 41, 51]. One meta-analysis examining case reports of human xylazine overdose found that in nonlethal cases where respiratory depression was observed and xylazine doses were quantified, doses ranged from 0.73 – 22 mg/kg [42]. These reports closely mirror our observation that doses of xylazine 1 mg/kg and above induce respiratory depression in mice. Medical literature is replete with case studies in which intravenous naloxone was administered to no apparent effect [29, 30, 35, 52–55]. Our results recapitulate this pattern and align with human clinical reports indicating that naloxone is not an effective reversal agent in cases of xylazine overdose.

In our studies, xylazine-adulterated fentanyl was not reversed by subsequent application of naloxone. A dose of naloxone that completely reversed the respiratory effects of fentanyl did not significantly reverse the respiratory suppressive effects of the same dose of fentanyl mixed with xylazine. Only two drugs – naloxone and nalmefene – are approved by the United States Federal Drug Administration for countering the respiratory effects of drug-induced overdose [56–59]. Both drugs function as competitive antagonists at MOR. However, we found that antagonism by naloxone is ineffective in reversing xylazine-involved respiratory depression. It is likely that only the opioid component of xylazine-adulterated fentanyl is affected by administration of naloxone. As no antidote [60] specific to xylazine-induced respiratory depression currently exists for human overdose, it is not clear whether any effective pharmacological tools exist to reverse overdoses inflicted by opioids adulterated by xylazine.

It has been suggested [61, 62] that atipamezole may serve as one such tool. The ability of atipamezole to reverse the respiratory effects of xylazine in humans has, to our knowledge, not been demonstrated. However, atipamezole effectively reverses the sedative and cardiovascular effects of α-2 adrenergic receptor agonists in humans [63] and α-2 antagonists have been shown to reverse the respiratory effects of MOR agonists in rats [64]. In our studies, atipamezole blocked xylazine’s suppression of breathing, and partially rescued fentanyl-induced respiratory depression in the absence of xylazine. In mice exposed to both xylazine and fentanyl, a subsequent injection containing a combination of atipamezole and naloxone produced rapid and complete reversal of respiratory suppression. In these mice, minute ventilation was restored to levels observed in mice exposed to vehicle within 25 minutes of naloxone-atipamezole administration, suggesting that combinations of naloxone and atipamezole may effectively reverse the respiratory effects of xylazine-fentanyl coadministration in cases of human overdose.

In contrast to naloxone, which rarely causes serious adverse effects in human clinical use [65, 66], atipamezole is known to increase arterial blood pressure and heart rate in humans, although it appears to be well-tolerated at the doses analogous to those required in our experiments to fully reverse the effects of xylazine-fentanyl coadministration [67–69]. Nevertheless, precipitated opioid withdrawal can also induce cardiovascular effects, which may impact the safety profile of interventions involving both naloxone and atipamezole [66, 70, 71]. While the potential benefits of therapeutic use of atipamezole in suspected cases of xylazine overdose may outweigh these risks, given the extensive and lethal nature of human use of adulterated illicit opioids and lack of existing reversal agents, caution must be exercised in interpreting these results. While we observed no fatalities in experimental animals, we did not evaluate the cardiovascular effects of atipamezole-naloxone coadministration in our experiments, and to our knowledge the potential cardiovascular interactions of naloxone-atipamezole coadministration remains unaddressed in current literature.

Functional interactions between MOR and α-2 receptors are supported in existing literature. Synergy and allosteric modulation between the analgesic effects of receptor agonism has been demonstrated in analgesia and antinociception [72, 73], and the receptors have been shown to dimerize with functionally interactions in cell culture [74]. Several canonical α-2 agonists demonstrate affinity for various opioid receptors [15, 75, 76]. α-2 receptors are co-expressed with MORs in the nucleus tractus solitarius [77], and play a functional role in the regulation of breathing by the retrotrapezoid nucleus [78] and locus coeruleus [79]. The receptors share a common signaling pathway, both coupling to Gi, and both interact with G protein-coupled inwardly rectifying potassium channels [80, 81], a key downstream target of MOR activation implicated in opioid-induced respiratory depression [82–84], though evidence for this effect is mixed *in vivo* [85]. Viewed holistically, these interactions suggest several potential pathways for the development of pharmacological interventions that simultaneously target respiratory depression induced by both drug classes. Respiratory stimulants agnostic to either receptor may also represent an effective strategy in combatting polydrug overdose resistant to reversal by naloxone [86], and this approach has shown promise in reversing CNS hypoxia inflicted by xylazine-fentanyl coadministration [87]. More work is necessary to determine the co-localization of α-2 and MORs within respiratory circuits and the mechanisms underpinning interactions between the respiratory effects of the two receptors.

Xylazine alone has been shown to induce CNS hypoxia at doses tightly coupled to those that depress respiration in our studies [38]. However, doses of xylazine that induce respiratory depression in both our studies and human overdose cases are far lower than those shown to induce lethality in mice, and we did not observe lethality in any experimental animals despite significant suppression of respiratory activity. In studies examining the lethality of xylazine, deaths were not observed until doses surpassed 60 and 100 mg/kg, i.p. [27, 28]. It has been suggested [38] that the hypoxic effects of xylazine may be independent of respiratory depression and instead may result from cerebral vasoconstriction or other well-defined effects of α-2 receptor activity. Our results suggest that it is unlikely that xylazine-induced CNS hypoxia is independent of respiratory depression, given the close coupling of doses demonstrated to depress breathing in our studies and the doses demonstrated to reduce partial oxygen in the central nervous system, though more work is necessary to determine whether the vascular effects of xylazine play a role in promoting xylazine-induced lethality. Another possibility is that the sustained interruptions of breathing responsible for lethality, which we did not measure, may only appear at higher doses of fentanyl and xylazine than those examined in our experiments.

In conclusion, we used whole body plethysmography to show that xylazine induces severe respiratory depression at subanesthetic doses in awake mice. This effect was dependent on α-2 adrenergic receptors and was fully blocked by the coadministration of the α-2 adrenergic receptor antagonist atipamezole. We further show that combinations of xylazine with low doses of fentanyl produce severe respiratory depression beyond that produced by either drug alone. Respiratory suppression inflicted by xylazine-fentanyl coadministration was not reversed by subsequent naloxone administration. In contrast, combined injection of naloxone and atipamezole fully reversed the respiratory effects xylazine-fentanyl coadministration. Taken holistically, these results suggest that xylazine may enhance lethality in humans by contributing to the respiratory suppression observed in opioid overdose, and that existing pharmacological tools to reverse lethal overdose are ineffective in cases involving xylazine.

## Acknowledgements

This work was supported by National Institutes of Health National Institute on Drug Abuse (NIDA) grant DA009158 (to AGH), DA047858 (to AGH), and Sigma Xi Grants-in-Aid of Research award G20240315-9075 (to JW). Whole body plethysmography equipment was funded in part with support from the Indiana Clinical and Translational Sciences Institute Research Equipment Fund ULTR002529 (to AGH). RH is supported by a Sidney and Becca Fleischer Research Scholarship. JW is supported by NIDA T32 Predoctoral Training Grant DA024628.

